# Phase-locking of resting-state brain networks with the gastric basal electrical rhythm

**DOI:** 10.1101/2020.10.06.328054

**Authors:** Ann S. Choe, Bohao Tang, Kimberly R. Smith, Hamed Honari, Martin A. Lindquist, Brian S. Caffo, James J. Pekar

## Abstract

A network of myenteric interstitial cells of Cajal in the corpus of the stomach serves as its “pacemaker”, continuously generating a *ca.* 0.05 Hz electrical slow wave, which is transmitted to the brain chiefly by vagal afferents. A recent study combining resting-state functional MRI (rsfMRI) with concurrent surface electrogastrography (EGG), with cutaneous electrodes placed on the epigastrium, found 12 brain regions with activity that was significantly phase-locked with this gastric basal electrical rhythm. Therefore, we asked whether fluctuations in brain resting state networks (RSNs), estimated using a spatial independent component analysis (ICA) approach, might be synchronized with the stomach. In the present study, in order to determine whether any RSNs are phase-locked with the gastric rhythm, an individual participant underwent 22 scanning sessions; in each, two 15-minute runs of concurrent EGG and rsfMRI data were acquired. EGG data from three sessions had weak gastric signals and were excluded; the other 19 sessions yielded a total of 9.5 hours of data. The rsfMRI data were analyzed using group ICA; RSN time courses were estimated using dual regression; for each run, the phase-locking value (PLV) was computed between each RSN and the gastric signal. To assess statistical significance, PLVs from all pairs of “mismatched” data (EGG and rsfMRI data acquired on different days) were used as surrogate data to generate a null distribution for each RSN. Of a total of 18 RSNs, three were found to be significantly phase-locked with the basal gastric rhythm, namely, a cerebellar network, a dorsal somatosensory-motor network, and a default mode network. Disruptions to the gut-brain axis, which sustains interoceptive feedback between the central nervous system and the viscera, are thought to be involved in various disorders; manifestation of the infra-slow rhythm of the stomach in brain rsfMRI data could be useful for studies in clinical populations.

## 1. Introduction

A network of myenteric interstitial cells of Cajal in the corpus of the stomach serve as its “pacemaker”, constantly and intrinsically generating a *ca.* 0.05 Hz electrical slow wave, which governs gastric peristalsis when there is food or chyme in the stomach, and which is transmitted to the brain chiefly by vagal afferents [1–4]. A recent study Rebollo et al. [5] combining resting-state functional MRI (rsfMRI) with concurrent surface electrogastrography (EGG), in which signals are recorded from cutaneous electrodes on the epigastrium (abdominal skin over the stomach), reported that brain activity in 12 regions including somato-motor cortices, dorsal precuneus, and the extrastriate body area was significantly phase-locked to the basal gastric rhythm. This collection of 12 gastric-synchronized regions, or nodes, was dubbed the gastric network, and it was suggested that time lags of several seconds between nodes were responsible for this “delayed connectivity network” not having been previously detected. This finding suggests that activity in brain resting-state networks (RSNs), estimated using network source separation techniques, such as the well-established spatial independent component analysis (ICA) approach, could be partially synchronized with the stomach. To ascertain whether any brain RSNs are synchronized with the gastric rhythm, we conducted a highly-sampled study in a participant who underwent 22 sessions of concurrent EGG and rsfMRI data collection, with two 15-minute runs per session, over a period of seven weeks. The resulting rsfMRI data were analyzed using spatial ICA to yield 18 RSNs, whose time courses were then tested for phase-locking with the basal gastric rhythm as determined from the concurrent EGG data. Three RSNs were found to be significantly phase-locked to the basal gastric rhythm, namely, a cerebellar network (FDR-adjusted p-value = 0.0022), a dorsal somatosensory-motor network (adjusted p-value = 0.0227), and a default mode network (adjusted p-value = 0.0227).

### 1.1. Resting-state functional MRI

Resting-state fMRI is a noninvasive neuroimaging method that uses MRI acquisitions originally developed to monitor hemodynamic sequelae of task-evoked changes in neuronal activity to observe neuronal activity in the brain “at rest” [6–9]. The resulting fMRI data manifest what are generally regarded as spontaneous fluctuations in intrinsic brain networks, allowing study of brain functional connectivity [10]. This methodology is popular not only because such data are easy to acquire, but also because they yield insights into a variety of conditions [7–9, 11–13]. For example, we have used rsfMRI to study patients with spinal cord injury, where paralysis could interfere with performance of motor tasks [14]. However, an important limitation of rsfMRI is that inter-regional synchrony of MRI time courses can result not just from synchronous neural events, but also from a variety of physiological sources [15] including cardiac pulsations [16, 17], respiration [18–20], and head motion [21, 22].

### 1.2. Resting-state brain networks

Resting-state fMRI originated from the observation that when the motor cortex peak voxel — the location with the highest fMRI activity during a motor task — was used as a seed to compute a map of temporal correlations from data acquired during rest, that the resulting resting-state network strongly resembled the motor task-activation map [6].

Independent component analysis is an exploratory data analysis approach that attempts to recover statistically independent sources from signals (data) that are modeled as mixtures of those sources [23]. The application of spatial ICA to fMRI data is broadly justified by the neurobiological principle of modularity, or the idea that different parts of the brain do different things. ICA was first applied to task fMRI data [24, 25], and then to rsfMRI [26–28], where it has become well-established [9, 13].

### 1.3. Visceral rhythms

The gut-brain axis, including neural, endocrine, and immune communication, is involved in the bidirectional interoceptive feedback between the central nervous system and the viscera. The brain monitors, and influences, the infra-slow rhythms generated in the viscera that control peristalsis. Even when the stomach is empty of food, electrical waves are constantly generated by myenteric interstitial cells of Cajal in the corpus of the stomach [1–4], with a normogastric period of about 20 seconds, or a frequency of approximately 0.05 Hz. Intestinal peristalsis is governed by ganglia of the enteric nervous system. These rhythms are communicated to the brain chiefly by the vagus nerve, and also the pelvic nerves of the parasympathetic nervous system, and the splanchnic nerves of the sympathetic nervous system.

### 1.4. Electrogastrography

Electrogastrography (EGG) uses cutaneous electrodes, placed on the epigastrium (abdominal skin lying above the stomach), in order to detect the gastric electrical slow wave [29–31]. Thus, EGG is similar to electrocardiography (ECG) and electroencephalography (EEG) in using surface electrodes to detect underlying bioelectrical signals. A chief difference is that EGG signals are much lower frequency than the corresponding signals from heart and brain, as the normogastric frequency in adults is approximately 0.05 Hz, or a period of 20 seconds. In the clinic, EGG is chiefly applied to patients with suspected motility disorders, such as indicated by recurrent episodes of nausea and vomiting. Recently there appears to be interest in applying EGG to psychophysiological research [32].

### 1.5. Concurrent rsfMRI and electrogastrography

A recent study Rebollo et al. [5] combined rsfMRI with concurrent EGG. Rebollo et al. [5] reported significant synchronization between the gastric rhythm and activity in a novel brain “gastric network” comprised of 12 nodes including somato-motor cortices, dorsal precuneus, and the extrastriate body area, with consistent inter-regional phase shifts or time lags. Because of these time lags, of several seconds between nodes, the novel gastric network was dubbed a delayed connectivity network, and the authors suggested that these delays were the reason that it had not been detected earlier using analytical approaches that look for inter-regional synchronization without such delays (but see [33]). An earlier report from the same group, using concurrent magnetoencephalography (MEG) and EGG, used a causal analysis to infer that the gastric rhythm was modulating regional cortical alpha-wave activity, presumably primarily via vagal afferent transmission [34].

### 1.6. Are any resting-state networks synchronized with the stomach?

Are any brain networks significantly phase-locked with the basal gastric rhythm? That is the question the present study addresses, in the context of brain resting state networks (RSNs) estimated using spatial ICA. To answer this question, we calculated the phase synchrony between each RSN time-course and concurrent EGG data using the phase-locking value (PLV). To assess statistical significance of these PLV values, we calculated PLVs for all pairs of “mismatched” data (EGG and rsfMRI data acquired on different days) to use as surrogate data in order to estimate the null distribution of PLV for each RSN. Comparing the matched and mismatched PLV distributions, we found that three brain networks were significantly phase-locked to the basal gastric rhythm: a cerebellar network (FDR-adjusted p-value = 0.0022), a dorsal somatosensory-motor network (adjusted p-value = 0.0227), and a default mode network (adjusted p-value = 0.0227).

## 2. Materials and methods

### 2.1. Experimental procedure

A healthy male volunteer, age 58, provided written informed consent to participate in a study approved by the Johns Hopkins Medicine Institutional Review Board. The participant was free of digestive, psychiatric, or neurological disorders, and had a body mass index (BMI) of 26. Twenty-two sessions were performed over a span of seven weeks. Data from three sessions were excluded due to weak gastric signals; data from the remaining 19 sessions were used for the analyses reported here. Scanning was typically performed on Mondays, Wednesdays, and Fridays. Sessions began at 9:00 AM; at 5:30 AM prior to each session, the subject breakfasted on multigrain cereal and yogurt, with coffee. The 3.5 hour period between breakfast and the session was intended to provide for gastric emptying [35]. The initial image acquisition was performed on 1 July 2019, and the last image acquisition was performed on 15 August 2019. Session dates are reported in Table 1.

**Table 1.** Session dates. Excluded sessions are crossed out using strikethrough.

|  | Mon | Tue | Wed | Thu | Fri |
| --- | --- | --- | --- | --- | --- |
| Week 1 | ~~7/1/2019~~ |  | 7/3 |  |  |
| Week 2 | 7/8 |  |  |  | 7/12 |
| Week 3 |  |  | 7/17 |  | ~~7/19~~ |
| Week 4 | 7/22 |  | 7/24 | 7/25 | 7/26 |
| Week 5 | 7/29 | 7/30 | 7/31 |  | 8/2 |
| Week 6 | 8/5 |  | 8/7 | 8/8 | 8/9 |
| Week 7 | 8/12 | ~~8/13~~ | 8/14 | 8/15 |  |

### 2.2. Magnetic resonance imaging

MRI was performed using a 3 Tesla Philips dStream Ingenia Elition scanner, operating at 45 mT/m with a slew rate of 200 mT/m/s. A multi-slice SENSE-EPI pulse sequence [36, 37] was used to acquire two resting state fMRI (rsfMRI) runs during each scanning session; the participant remained in the scanner between runs. Each run was acquired using the following acquisition parameters: acquisition time = 15 *min*, TR/TE = 2000/30 *ms*, number of dynamics = 450, field of view = 240 × 240 *mm*^2^, 3-*mm* isotropic spatial resolution, 36 axial slices collected sequentially in increasing slice order with a 1-*mm* gap, SENSE acceleration factor = 2, and flip angle = 71°. Respiratory rate was simultaneously measured using a pulse oximeter. The participant was instructed to rest comfortably while remaining still, and no other instruction was provided. The subject’s eyes were closed for the rsfMRI acquisitions. A T1-weighted (T1w) Magnetization-Prepared Rapid Acquisition Gradient Echo (MPRAGE) structural run was acquired during the third session (on July 12) using the following acquisition parameters: acquisition time = 5 *min*, TR/TE/TI = 10/6/842 *ms*, field of view = 212 × 212 *mm*^2^, resolution = 1.1 × 1.1 × 1.2 *mm*^3^, 120 axial slices collected, SENSE acceleration factor = 2, and flip angle = 8°). The T1w images were subsequently used to align and normalize the fMRI images.

### 2.3. Electrogastrography

The gastric rhythm of the participant was monitored using MRI compatible electrogastrography (EGG) equipment (BIOPAC MP160 system; BIOPAC Systems Inc, USA). Acquisition parameters and placement of cutaneous electrodes, similar to those described by Rebollo et al. [5], are summarized here.

For preparation, intended electrode locations were marked on the participant’s epigastrium (see Fig 1(a)), then the marked regions were rubbed and cleaned with alcohol to remove dead skin, and electrolyte gel was applied. Three sets of bipolar electrodes were then placed in two rows over the abdomen. EGG was then acquired at a sampling rate of 200 Hz with a low-pass filter of 1 Hz and no high-pass filter.

**Figure 1.**
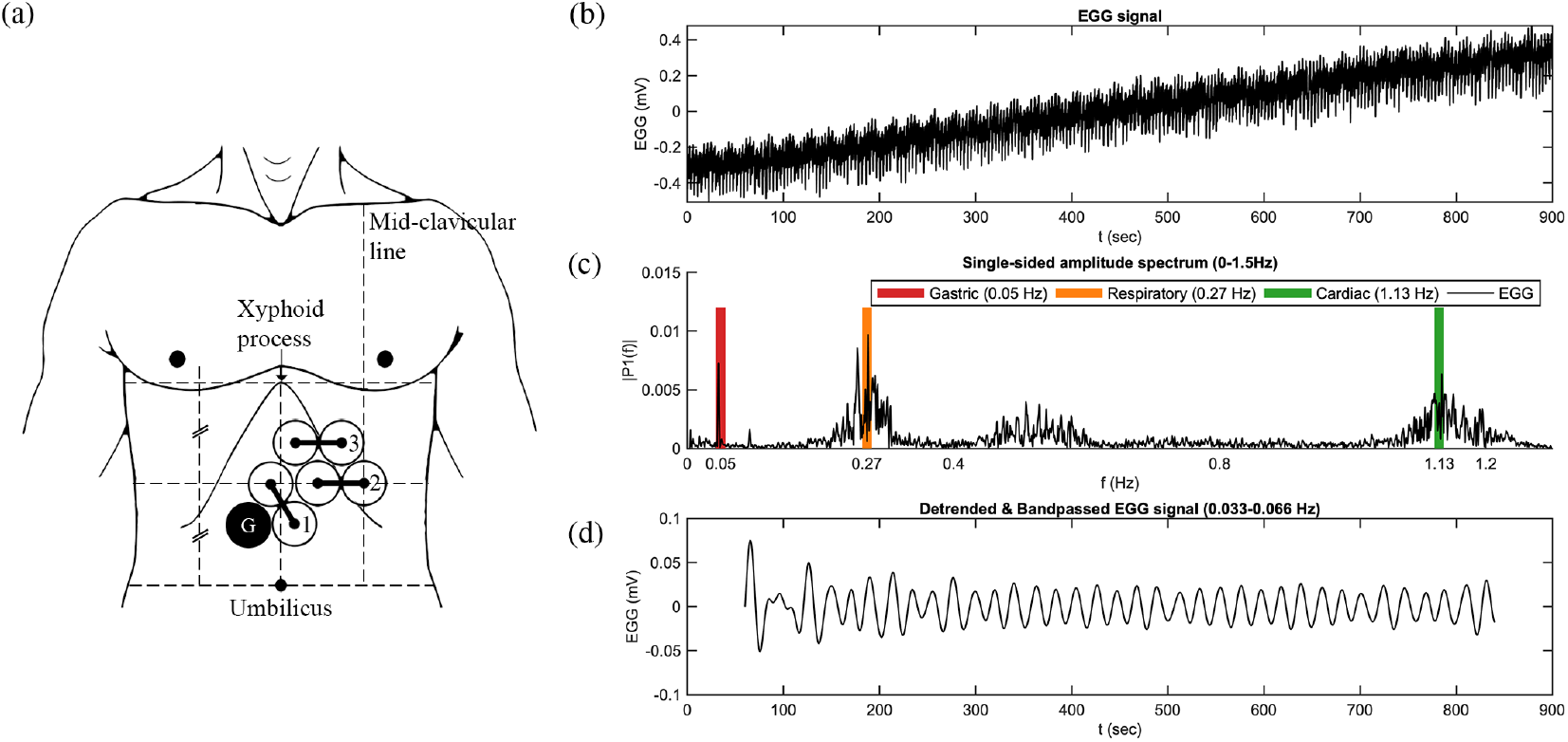
Electrogastrography electrode placement and representative data. (a) Electrode placement. (b) Representative EGG signal time course. (c) Representative signal spectrum (Fourier transform of the detrended EGG signal shown in part (b)). (d) Gastric signal after detrending and bandpass filtering.

Prior to the acquisition of any MRI data, five minutes of EGG data were acquired with the participant lying outside the tunnel of the scanner. This was to acquire a reference EGG signal with a frequency content free of the effect of the static magnetic field and gradient pulses. Once the participant was placed inside the scanner, EGG data were recorded concurrently with rsfMRI data.

### 2.4. Data analysis

#### 2.4.1. RsfMRI preprocessing

Preprocessing of the rsfMRI data set was performed using the Analysis of Functional NeuroImages (AFNI) software (version AFNI 20.1.06) [38]. The preprocessing pipeline included: 1) despiking, 2) slice timing correction, 3) motion correction, 4) co-registration, 5) normalization, 6) segmentation, and 7) spatial smoothing using a 6 mm (i.e., twice the nominal acquisition voxel size) full-width at half-maximum Gaussian kernel.

#### 2.4.2. RsfMRI independent component analysis

The Group ICA of fMRI Toolbox (GIFT) software (http://trendscenter.org/software/gift/; version v4.0b) [39] was used to perform group independent component analysis (GICA) [40]. Two steps of principal component analysis (PCA) data reduction were performed for group level analysis, where individual session data were first reduced to 84 principal components. The reduced data were then concatenated in the temporal direction and further reduced to 42 principal components. Estimation of the number of independent components (i.e., 42) was guided by order selection using the minimum description length (MDL) criterion [41]. The dimensionality of the individual session PCA data reduction (i.e., 84) was set by doubling the estimated component number, to ensure robust backreconstruction [42, 43].

The spatial distribution (i.e., grey matter vs. white matter and cerebral spinal fluid) and temporal frequency power distribution of 42 ICs were manually assessed using the aggregate spatial maps and time courses, and 22 ICs were eliminated as representing non-neuronal sources such as head motion, respiration, and cardiac pulsations. Two additional ICs were rejected due to low similarity measures calculated using the ICASSO toolbox [44]. This process identified the remaining 18 ICs as functional RSNs, which are shown in Fig 2.

**Figure 2.**
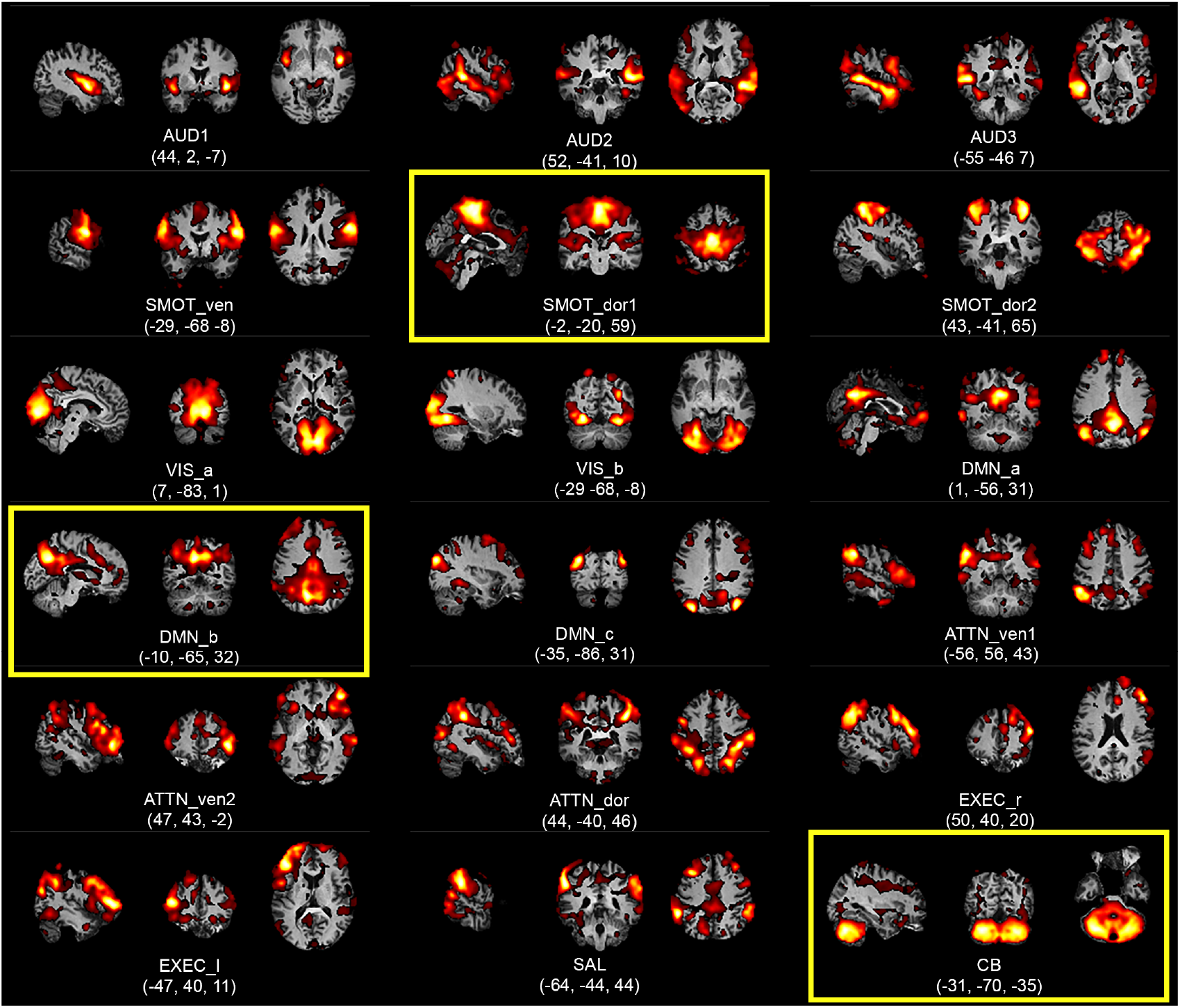
Aggregate spatial maps of the resting state networks (RSNs) of the highly sampled participant. Group independent component analysis (GICA) was used to estimate the RSNs and obtain the aggregate spatial maps. The spatial maps of each RSN are shown as subfigures, with representative sagittal, coronal, and axial views (left-to-right) overlaid on structural images within the Montreal Neurological Institute (MNI) template space; three RSNs with statistically significant gastric phase-locking are highlighted using yellow boxes. Coordinates (in mm) for each view are indicated below each subfigure. (AUD: auditory, SMOT: somatosensory-motor, VIS: visual, DMN: default mode network, ATTN: attention, EXEC: executive, SAL: salience, CB: cerebellar, ven: ventral, dor: dorsal, r: right, l: left).

Single-session maps and time courses for each session were obtained using “GICA1” back-reconstruction [42, 45].

#### 2.4.3. EGG preprocessing

EGG preprocessing was performed following the pipeline developed by Rebollo et al., [5], using the FieldTrip toolbox (http://www.fieldtriptoolbox.org/) [46], Matlab (Natick, MA; version R2018a), and custom code provided by Rebollo et al. [5] (https://github.com/irebollo/stomach_brain_Scripts). Data were low-pass filtered below 5 Hz to avoid aliasing of higher-frequency signals, e.g., cardiac, and down-sampled to 10 Hz. To identify the EGG peak frequency (0.033–0.066 Hz) for each run, we computed the spectral density estimate for each EGG channel over the 900 s of EGG signal acquired during each fMRI scan using Welch’s method on 200 s time windows with 150 s overlap. For each run, the spectral peak was identified by looking for a sharp peak within the normogastric frequency range of 0.033–0.066 Hz. Data from the EGG channel with the highest spectral peak were then bandpass filtered to isolate the signal related to gastric basal rhythm (linear phase finite impulse response filter, FIR, designed with Matlab function FIR2, centered at EGG peaking frequency, filter width ± 0.015 Hz, filter order of 5). Data were filtered in the forward and backward directions to avoid phase distortions and then further downsampled to match the sampling rate of the BOLD acquisition (0.5 Hz).

#### 2.4.4. Quantification of RSN–EGG synchronization

The RSN time courses were bandpass-filtered using the same filter parameters that had been applied to the EGG data from the corresponding run. To avoid edge effects, the first and last 15 volumes (30 s) were discarded from both the RSN and EGG time courses. The updated duration of the fMRI and EGG signals for which the rest of the analysis was performed was thus 840 s. The Hilbert transform was applied to the filtered RSN and EGG time courses to derive their instantaneous phases. The phase-locking value (PLV) [47] was computed as the absolute value of the time average difference in the angle between the phases of the EGG and the fMRI time course (Equation 1).

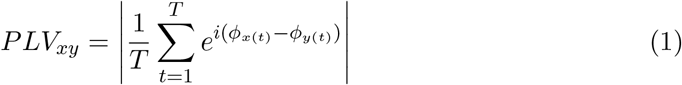

where x and y are the two time series, and T is the number of samples in each time series. The PLV ranges between the values of 0 and 1, where 0 represents no synchrony, and 1 represents perfect synchrony. Notably, the PLV measures are independent of temporal delays and amplitude fluctuations of the input signals. The PLV was averaged over the whole duration of the recording.

#### 2.4.5. Statistical analysis of RSN–EGG synchronization

We implemented a two-step statistical procedure, based on methodology used by Rebollo et al. [5]. We first estimated chance-level phase-locking between brain networks and the stomach, i.e., PLV expected in the absence of true stomach-RSN synchronization. Then we applied a multiple testing procedure to identify those RSNs in which measured gastric-brain phase-locking was significantly greater than chance. Here, chance phase-locking was defined via negative control comparisons based on the observed data.

Specifically, surrogate data were used to estimate null distributions representing chance-level PLV. We used mismatched data, i.e., all pairs of EGG and rsfMRI data acquired on different days. For each RSN, surrogate data were created by calculating the PLV between that RSN time course and all gastric time courses that were acquired on different days. As we used data from two runs each on 19 days, this mismatching approach yields 1,368 surrogate data entries representing a null distribution for PLV for each RSN.

Comparisons of the observed PLVs to this null distribution yield information on how likely results were compared to chance-level PLVs. Therefore, in the second step, we formally tested whether, for each RSN, the empirical PLV differed from chance-level PLV. For each RSN, we applied a Wilcoxon rank test (since PLVs are clearly non-Gaussian) to test whether there was a significant mean difference between empirical PLVs from 38 runs against the surrogate data. To correct for multiple comparisons, the p-values for the 18 RSNs were adjusted using the Benjamini-Hochburg procedure [48] with a false discovery rate (FDR) of 0.05, in order to judge which RSNs were significantly phase-locked with the gastric rhythm.

#### 2.4.6. Phase-locking of gastric signals between different runs

To shed light on the variability of the subject’s gastric rhythm, we computed the PLV of the gastric rhythm from different runs. Specifically, we compared the distribution of PLVs computed from runs from the same day, to the distribution of PLVs computed from runs from different days. We then used the Wilcoxon rank test to compare the same-day and different-day gastric-gastric PLV distributions.

#### 2.4.7. Quantification of gastric contributions to rsfMRI signals

To estimate the magnitude of the manifestation of the infra-slow gastric rhythm in RSN time courses, we calculated the proportion of the rsfMRI signal variance, or perecent variance accounted for (P.V.A.F), that could be explained by the EGG signal. P.V.A.F here is represented by the R-squared *R*^2^ of the corresponding linear regression after adjusting for any phase delays between the two signals.

First, we assumed that the instantaneous phase difference Δ*ψ* = *ψ*(EGG) − *ψ*(rsfMRI) i.i.d. follows a von Mises distribution *p*(Δ*ψ*|*μ, κ*) ∝ exp(*κ* cos(Δ*ψ* − *μ*)), which is a typical circular distribution used to model phase differences [49]. Then we performed maximum likelihood estimation for the positional parameter *μ* as the over-all phase difference between two signals.

Second, we adjusted such phase difference by multiplying the Hilbert transformation of the rsfMRI signal by 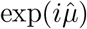 for 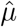 the MLE of *μ* above. Then we calculated as the P.V.A.F. the *R*^2^ between EGG signal regressing on the real part of this new signal. This definition of P.V.A.F takes phase-locking into consideration and makes any necessary adjustment to align signals before regression.

## 3. Results

We found 18 RSNs, as shown in Fig 2.

Electrode pair 2 (as illustrated in Fig 1(a)) consistently gave the best gastric signals. The subject’s gastric rhythm was within the normogastric range, at 0.048 ± 0.001 Hz.

For each RSN, we calculated its PLV with respect to gastric phase for each scan, and also the similar PLV, but for mismatched data pairs acquired on different days. For each RSN, we compared the distribution of measured PLVs with the distribution of mismatched PLVs using the Wilcoxon rank test. These results are shown in Fig 3 and tabulated in Table 2. The table gives the uncorrected p-values as well as p-values adjusted for multiple comparisons using the Benjamini-Hochberg method [48] for a FDR of 0.05.

**Figure 3.**
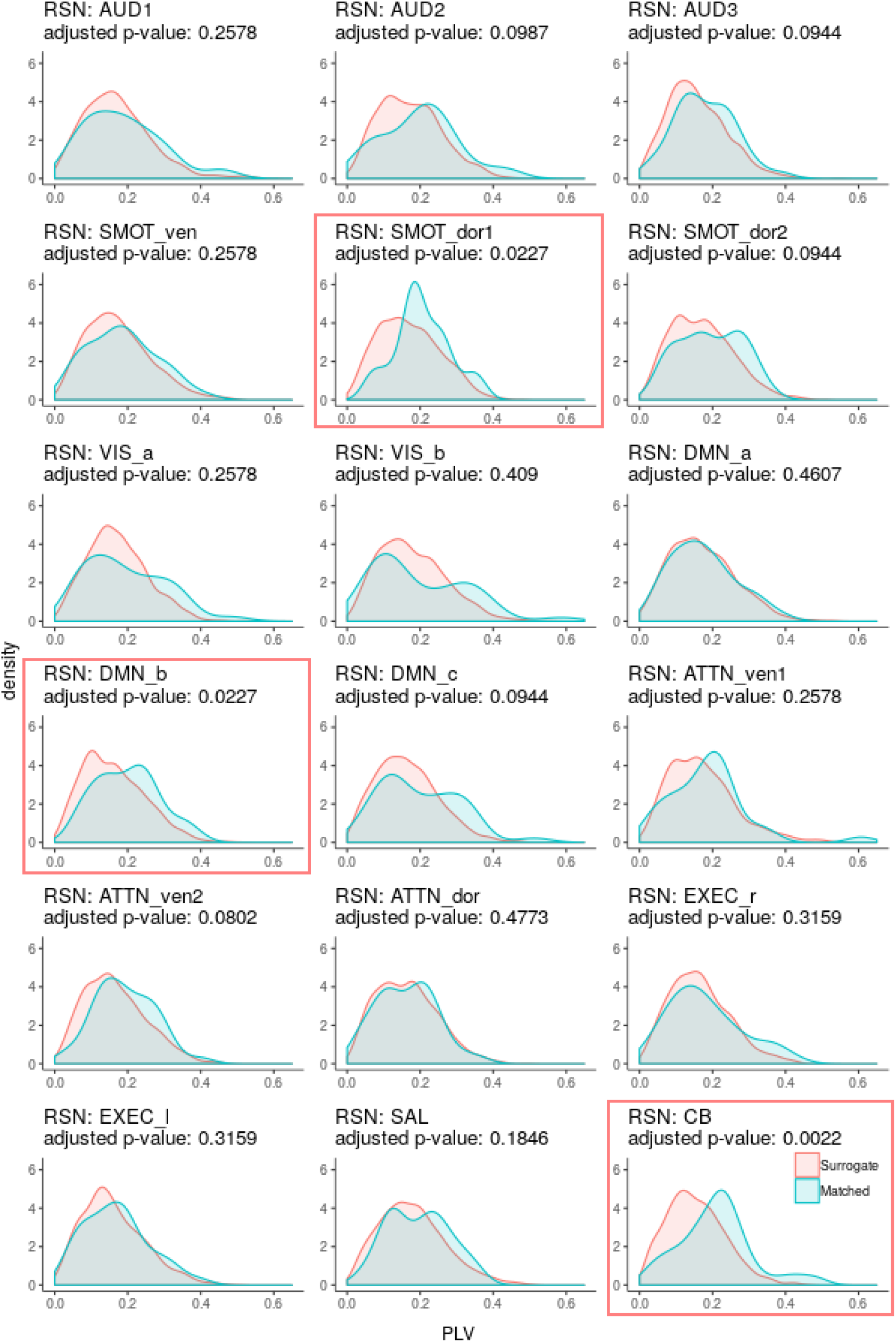
Gaussian kernel density estimates (i.e., smoothed histograms) of PLV between resting-state networks and gastric signals recorded concurrently (cyan) and on different days (coral). P-values from Wilcoxon rank tests have been adjusted for multiple comparisons using a FDR of 0.05.

**Table 2.** Phase locking value (PLV) between resting state networks (RSNs) and electrogastrography (EGG) signals.

| RSN | PLV mean | PLV std. dev. | Mismatched PLV mean | Mismatched PLV std. dev. | Uncorrected p-value | FDR-adjusted p-value | Fractional gastric variance |
| --- | --- | --- | --- | --- | --- | --- | --- |
| AUD1 | 0.1882 | 0.1056 | 0.1709 | 0.0935 | 0.1852 | 0.2578 | 0.0111 |
| AUD2 | 0.1995 | 0.1040 | 0.1725 | 0.0860 | 0.0439 | 0.0987 | 0.0266 |
| AUD3 | 0.1791 | 0.0806 | 0.1567 | 0.0808 | 0.0330 | 0.0944 | 0.0064 |
| SMOT_ven | 0.1854 | 0.0941 | 0.1729 | 0.0891 | 0.1862 | 0.2578 | 0.0038 |
| SMOT_dor1 | 0.2061 | 0.0758 | 0.1697 | 0.0845 | 0.0029 | 0.0227 | 0.0150 |
| SMOT_dor2 | 0.1947 | 0.0855 | 0.1705 | 0.0855 | 0.0367 | 0.0944 | 0.0057 |
| VIS_a | 0.1939 | 0.1078 | 0.1711 | 0.0815 | 0.1619 | 0.2578 | 0.0338 |
| VIS_b | 0.1978 | 0.1288 | 0.1775 | 0.0911 | 0.3636 | 0.4090 | 0.0420 |
| DMN_a | 0.1746 | 0.0882 | 0.1719 | 0.0892 | 0.4351 | 0.4607 | 0.0242 |
| DMN_b | 0.1990 | 0.0857 | 0.1612 | 0.0856 | 0.0038 | 0.0227 | 0.0291 |
| DMN_c | 0.1987 | 0.1085 | 0.1639 | 0.0849 | 0.0346 | 0.0944 | 0.0185 |
| ATTN_ven1 | 0.1860 | 0.1078 | 0.1705 | 0.0920 | 0.1504 | 0.2578 | 0.0071 |
| ATTN_ven2 | 0.1920 | 0.0812 | 0.1651 | 0.0845 | 0.0178 | 0.0802 | 0.0139 |
| ATTN_dor | 0.1620 | 0.0804 | 0.1634 | 0.0834 | 0.4773 | 0.4773 | 0.0201 |
| EXEC_r | 0.1797 | 0.0996 | 0.1645 | 0.0822 | 0.2523 | 0.3159 | 0.0071 |
| EXEC_l | 0.1689 | 0.0885 | 0.1603 | 0.0854 | 0.2633 | 0.3159 | 0.0079 |
| SAL | 0.1952 | 0.0831 | 0.1785 | 0.0902 | 0.0923 | 0.1846 | 0.0095 |
| CB | 0.2093 | 0.1002 | 0.1539 | 0.0791 | 0.0001 | 0.0022 | 0.1124 |

AUD: auditory, SMOT: somatosensory-motor, VIS: visual, DMN: default mode network, ATTN: attention, EXEC: executive, SAL: salience, CB: cerebellar ven: ventral, dor: dorsal, r: right, l:left std. dev.: standard deviation, FDR: false discovery rate

Three networks were significantly phase-locked with the basal gastric rhythm: A cerebellar network (CB; adjusted p-value = 0.0022), a dorsal somatosensory-motor network (SMOT dor1; adjusted p-value = 0.0227), and a default mode network (DMN b; adjusted p-value = 0.0227). The fraction of (gastric band-passed) RSN signal variance that could be accounted for by the gastric rhythm, using linear models, was about 1.5 percent and three percent for the two cortical RSNs that were significantly phase-locked with the gastric basal electrical rhythm, and about 11 percent for the cerebellar network.

Distributions of phase-locking values between pairs of gastric signals from different runs are presented in Fig 4.

**Figure 4.**
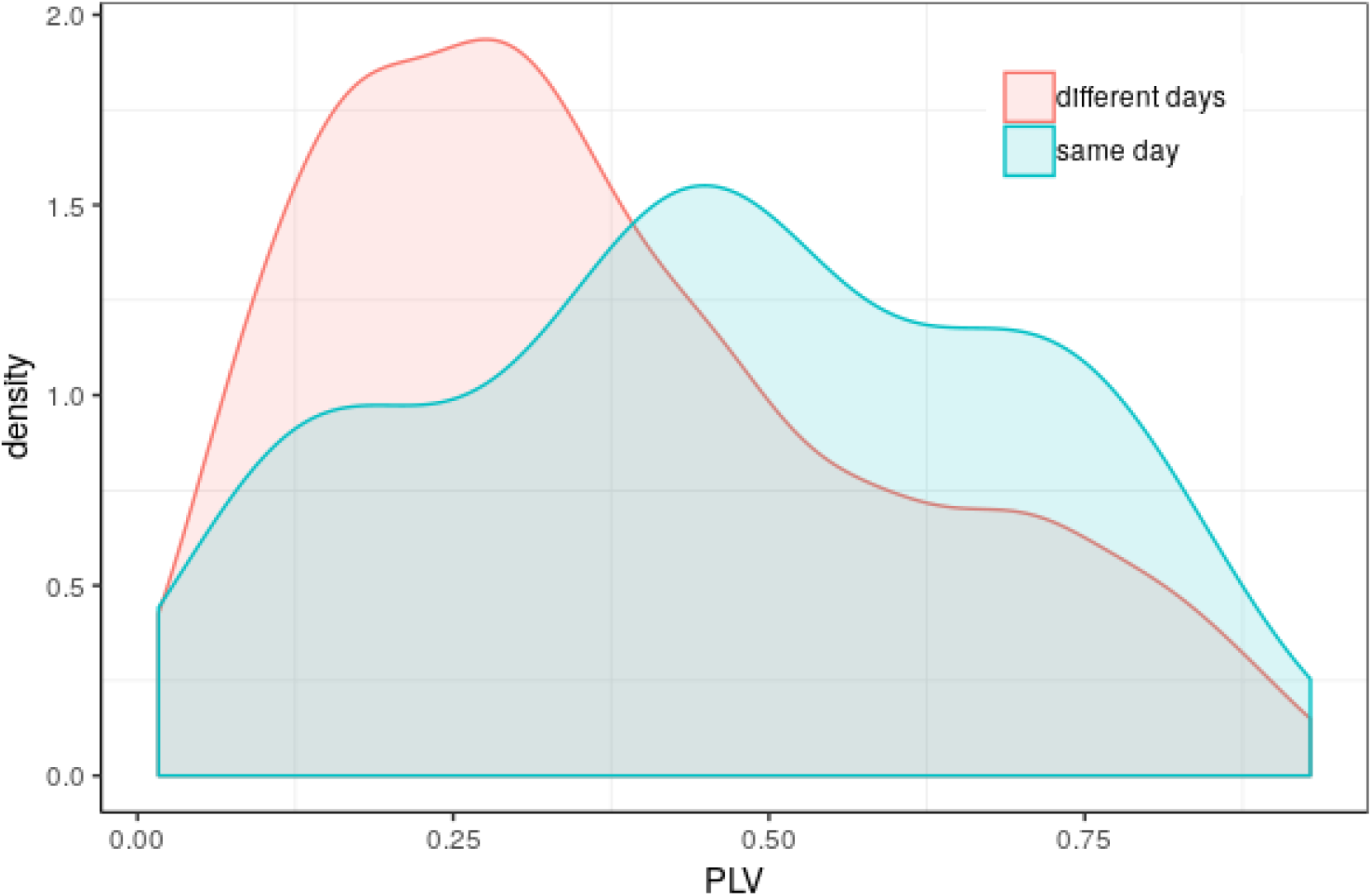
Gaussian kernel density estimates of PLV for pairs of gastric signals from the same day (cyan) and different days (coral). The Wilcoxon rank test p-value was 0.00665.

## 4. Discussion

In this study, eleven hours of concurrent resting state fMRI / surface electrogastrography (EGG) data were acquired in 22 sessions over seven weeks. Three sessions resulted in EGG data with weak gastric signals and were excluded; the 9.5 hours of data from the remaining 19 sessions were analyzed using spatial ICA, yielding 18 resting state brain networks (RSNs). Three of the RSNs were then found to be significantly phase-locked with the basal gastric rhythm as estimated from the EGG data, namely, a cerebellar network (FDR-adjusted p-value = 0.0022), a dorsal somatosensory-motor network (adjusted p-value = 0.0227), and a default mode network (adjusted p-value = 0.0227).

### 4.1. What is your stomach saying to your brain?

Evidence that rsfMRI signals are widely regarded as spontaneous is provided by a Google scholar search for “’spontaneous fluctuations’ and rsfMRI”, which returns over 5,000 citations (as of June 2020). In light of the findings of Rebollo et al. [5], and of the present study, it may be useful to examine the concept of spontaneity in the context of resting-state functional neuroimaging. The sense of “spontaneous” that applies here appears to be the third definition given by the Oxford English Dictionary [50]: “Of natural processes: Occurring without apparent external cause; having a self-contained cause or origin.” These fluctuations are indeed spontaneous; the context in which they are self-contained must include the stomach, and presumably other organs of the body, as well. This is consistent with the principle of interoceptive cognition, the view that the brain’s home in the body is fundamental to its function [51–53], or that, to quote the title of a recent review: “Visceral signals shape brain dynamics and cognition” [54]. In other words, by enlarging our scope beyond the central nervous system and considering the entire (organismic) nervous system—including the enteric nervous system— brain fluctuations and the gastric rhythm are similarly seen as spontaneous manifestations of intrinsic networks.

In the present study, an effort was made to establish a similar gastric state at each experimental session, using a fixed schedule and fixed breakfast. Researchers have probed the effect of gastric state, including overnight fasting, on rsfMRI outcomes [55, 56]. Concurrent EGG would allow future studies in this area to benefit from measures of how gastric state affects stomach-brain synchronization.

In the present study, we acquired fMRI data only in the “resting” state. Researchers have used fMRI to measure brain responses to hedonic/gustatory stimuli such as photographs of food [57] and the the ingestion of small samples (“sips”) of fluid delivered during scanning [58, 59]. Concurrent EGG would allow future studies in this area to benefit from measures of how such stimuli affect the stomach’s rhythm, and stomach-brain synchronization.

How much data per person would be needed for concurrent fMRI/EGG studies in clinical populations? The RSN-stomach PLV values in Table 2 and Fig 3 can be interpreted as providing an estimate of the precision of such measures, in terms of mean and standard deviation PLV. It appears that the relative precision of our significant phase-locking estimates that would be derived from a single 15-minute scan, would be on the order of plus-or-minus 30 percent, which is relatively modest. However, we do not know how the inter-session variability of other participants would compare to that of ours; similarly, we cannot speculate about the inter-individual variability of these measures in various populations of interest.

### 4.2. Measuring synchronization using the phase-locking value

How should synchrony be quantified? A recent review lists no fewer than fifteen measures of “brain functional connectivity through phase coupling of neuronal oscillations” [60]. In the present study, we followed Rebollo et al. [5], in using the phase-locking value (PLV) to assess infra-slow stomach-brain synchrony. Here, the PLV has two major advantages: The PLV is insensitive to variations in amplitude, which is attractive because we are interested only in the synchrony of stomach and brain fluctuations, regardless of their amplitude. The PLV seeks only consistent phase offsets (it does not require zero phase offset), which is attractive because we do not know what lags may be present between stomach electrical activity and synchronized but hemodynamically-delayed brain BOLD signals.

### 4.3. Testing for significance of the phase-locking value

To test for the statistical significance of phase-locking, surrogate data were used in order to generate an estimate of the chance distribution of PLV, which would be obtained in the absence of true stomach-brain synchronization. To generate such surrogate data, we used all pairs of mismatched data, that is, rsfMRI and EGG data acquired on different days. This appears reasonable because one would not expect today’s brain to be synchronized with yesterday’s stomach. However, if the gastric basal electrical rhythm were like a tuning fork, always ringing at a never-changing frequency, then today’s gastric rhythm would be phase-locked to yesterday’s gastric rhythm. Hence, to the extent that the gastric rhythm may be unexpectedly stable, then our use of mismatched data as surrogate data would therefore be overly conservative. In fact, as shown in Fig 4, the subject’s gastric rhythm was sufficiently stable that his average within-day PLV between the two 15-minute gastric scans was larger than his average between-day PLV between gastric scans acquired on different days.

Rebollo et al. [5] used mismatched data as surrogate data, but theirs was a group study of 34 people, each scanned once. So for their study, mismatched data were data from other people, not from the same person on other days.

To correct for multiple comparisons, we used the Benjamini-Hochberg approach to adjust p-values using a False Discovery Rate of 0.05 [48]. Using instead the conservative Bonferroni approach to correcting p-values, then only the cerebellar network would be judged significantly phase-locked with the stomach; the dorsal somatosensory-motor network would have a Bonferroni corrected p-value of 0.053, just above the 0.05 threshold. However, that the Bonferroni correction is overly conservative when it comes to testing time courses derived using spatial ICA can be seen from the fact that such time courses are generally not independent; indeed, measures of inter-network temporal correlations are a subject of study [61, 62].

### 4.4. On gastric phase-locking of cerebellar, dorsal somatosensory-motor, and default mode networks

We estimated a total of 18 RSNs, which were broadly similar to those reported in earlier studies that applied ICA to rsfMRI data [9, 13, 26–28, 63]. We found that three of the 18 RSNs were phase-locked with the stomach. While the role played by gastric synchronization of brain networks is unknown, the gastric phase-locking of these three networks appears to be consistent with literature on their involvement in feeding behavior. In the case of the cerebellum, early evidence was provided by small-animal experiments, in which electrical stimulation of the cerebellum induced feeding behavior [64]; a range of evidence now provides support for the role of the cerebellum in eating [65]. The dorsal somatosensory-motor network includes the primary somatosensory cortex and the medial wall motor regions, two of the nodes reported by Rebollo et al. [5]; both of these regions contain body maps, presumably including representations of the stomach. The default mode network significantly phase-locked with the gastric basal electrical rhythm is centered on the precuneus, which was a node of the gastric network reported by Rebollo et al. [5], and has been reported to be involved in appetite control in healthy individuals [66], and its disruption in persons with obesity [66] as well as anorexia and bulemia [67].

### 4.5. On highly-sampled individual brains

This manuscript reports on a highly-sampled individual brain. The eleven hours of data we collected is perhaps 40 times as much data per brain as was often seen in the general task fMRI literature. The approach of acquiring much more data from many fewer brains appears to have been originally suggested by Savoy [68], and popularized by Poldrack [69] and a group at Washington University St. Louis [70]. We previously reported on data from an individual who underwent weekly scans over a multi-year period [63]. Highly-sampled data have also led to further analyses and publications, e.g., [71, 72].

### 4.6. Limitations

A major limitation of the present report is that the eleven hours of data were acquired from a single person, as part of a highly-sampled individual study design. Hence, it is not clear how well these results will generalize to the broader population, and it is of course necessary to scan more people in order to find out.

A limitation of the present report is that experimental procedures, including cutaneous electrodes on the epigastrium and mild fasting, may have focused the participant’s attention on his gastric state. However the participant reported typical wandering thoughts, without a sustained or noticeable focus on hunger.

A limitation of the present report is that by adhering to the same schedule and same breakfast, our data may be unusually stable with regards to variance over sessions. Data acquired “in the wild” will presumably manifest more variation.

A limitation of the present report is that we did not acquire any physiological measures (e.g., blood glucose) on a daily basis, in order to help explain inter-session variance. It might be advisable to collect such measures in future population-based studies.

A limitation of the present report is that, during scanning, we did not acquire other real-time physiological measures, such as the electrocardiogram. (Although cardiac events are seen in the raw electrogastrography data, they are in a distinct frequency range from the infra-slow basal gastric rhythm, and so do not alias into or contaminate the estimated gastric rhythm.) Future studies might monitor a comprehensive suite of physiological measures during “rest”, to shed light on activity fluctuations of the sympathetic and parasympathetic nervous systems. A group from Arhus University recently announced that they are acquiring rsfMRI data with concurrent “arterial CO2, respiratory, cardiac, eye tracking, and electrogastrography measures” [73].

A limitation of the present report is that we used only one analytic approach, spatial ICA, to estimate resting state brain networks. In keeping with the “plurality and resemblance” framework [74], it would be beneficial to re-analyze the data using alternative approaches (e.g., atlas-based parcellations or graph-theoretic methods) in order to understand how the results presented here resemble those obtained using other analytic approaches to estimating brain networks from rsfMRI data.

A limitation of the present report is that these data cannot be used to infer directionality or causality. Simply from the EGG and rsfMRI data alone, we cannot tell whether the stomach is driving the brain, or whether the brain is driving the stomach. The role of myenteric interstitial cells of Cajal as the generators or “pacemakers” of visceral rhythms, which are transmitted to the brain, appears firmly established [1–4]. In the context of resting-state functional neuroimaging, an earlier study from the same group at the École normale supérieure (Paris) that contributed the Rebollo et al. [5] paper, supports ascending directionality: Richter, et al. [34], used concurrent EGG and magnetoencephalography (MEG); the high time resolution of MEG allowed causal inference, which indicated that the gastric rhythm was modulating regional alpha-band brain activity. Rebollo et al. [5] concluded that their results supported “…the hypothesis that activity in the gastric network is driven by neural activity in areas directly receiving ascending inputs…” However, the brain is not simply a passive recipient of visceral rhythms. Rather, “[w]ithin the brain, adaptive control is achieved through forward models, efference copies and prediction errors wherein viscerosensory data is continuously compared against expected bodily state to evoke physiological and mental reactions” [51]. That is, it appears reasonable to surmise that the brain contains one or more interoceptive representations of the stomach, which model its rhythm, and that these dynamic models are entrained to the actual stomach rhythm, using prediction errors. Thus, when several brain regions are found with activity that is synchronized with the stomach, it may not be clear whether a particular region’s gastric-synchronized brain signal results from ascending afferent signals, or from an ongoing brain dynamic model of the stomach. At a very different time scale (and presumably subserved by very different mechanisms), an analogy with the (*ca.* 4000-fold slower) chronobiology of circadian rhythms [75] may be helpful here, as these diurnal rhythms are not simply driven by external stimuli, but rather are generated internally by “biological clocks”, which are entrained to the local astronomical day/night cycle. Finally, the success of electrogastrographic biofeedback [76], in which individuals provided with real-time visual feedback on their gastric rhythm are able to improve its consistency, demonstrates that the brain can influence the gastric rhythm, and there has been recent progress in understanding how cortical regions communicate with the stomach [77]. Clearly, further studies are needed to explore the bidrectional communication between the viscera and the brain.

### 4.7. A complementary view: Gastric synchronization as a confound

As mentioned, a fundamental limitation of the general rsfMRI approach is that a variety of “nuisances” or physiological confounds [15] including cardiac pulsations [16, 17], respiration [18–20], and head motion [21, 22], can lead to inter-regional correlations that can mask, or be mistaken for, functional connectivity. The manifestation of the infra-slow gastric rhythm in brain fMRI data could be seen as a similar confound, consistent with recent guidance that any “neural activity related fluctuations that are not of interest” shall be regarded as “physiological noise” [78]. Thus, if stomach-brain synchronization is not celebrated as a window onto the embodied brain in the organismic nervous system, but instead regretted as a nuisance — if the gastric rhythm is the new head motion — then the question arises as to its magnitude. The modest contributions tabulated in Table 2 — less than three percent for the two gastric phase-locked cortical networks, and about eleven percent for the cerebellar network — suggest that if the gastric rhythm is seen is a confound, it may not be a serious one.

## 5. Conclusion

Of 18 resting-state brain networks estimated from rsfMRI data using the well-established spatial ICA approach, three were found to be significantly phase-locked with the basal gastric rhythm, namely, a cerebellar network, a dorsal somatosensory-motor network, and a default mode network. Disruptions to the gut-brain axis, which sustains interoceptive feedback between the central nervous system and the viscera, are thought to be involved in various disorders; manifestation of the infra-slow rhythm of the stomach in brain rsfMRI data could be useful for studies in clinical populations.

## Acknowledgements

We are grateful to Peter A. Barker, Terri Brawner, Jiande Chen, Joseph S. Gillen, Kathleen Kahl, Ivana Kusevic, and Peter C.M. van Zijl for assistance and encouragement. Support was provided by the National Institutes of Health, USA, under grants NIH 5R21NS104644-02 (A.S.C.); NIH R01EB016061 (M.A.L.); NIH 1R21EB030009-01 (J.J.P.); NIH 1U54HD079123-03 (S.A. Fatemi); NIH 5P41EB015909-19 (P.C.M. van Zijl); and NIH 1S10OD021648 (P.C.M. van Zijl). The funders had no role in study design, data collection and analysis, decision to publish, or preparation of the manuscript.

